# The role of thickness inhomogeneities in hierarchical cortical folding

**DOI:** 10.1101/2020.04.01.020172

**Authors:** Lucas da Costa Campos, Raphael Hornung, Gerhard Gompper, Jens Elgeti, Svenja Caspers

**Affiliations:** Theoretical Physics of Living Matter, Institute of Biological Information Processing (IBI-5), Research Centre Jülich, Jülich, Germany; Institute of Neuroscience and Medicine (INM-1), Research Centre Jülich, Jülich, Germany; JARA-Brain, Jülich-Aachen Research Alliance, Jülich, Germany; Institute for Anatomy I, Medical Faculty, Heinrich-Heine University, Düsseldorf, Germany

**Keywords:** cortical folding, cortical thickness, gyrogenesis, gyrification, higher-order folding

## Abstract

The morphology of the mammalian brain cortex is highly folded. For long it has been known that specific patterns of folding are necessary for an optimally functioning brain. On the extremes, lissencephaly, a lack of folds in humans, and polymicrogyria, an overly folded brain, can lead to severe mental retardation, short life expectancy, epileptic seizures, and tetraplegia. The construction of a quantitative model on how and why these folds appear during the development of the brain is the first step in understanding the cause of these conditions. In recent years, there have been various attempts to understand and model the mechanisms of brain folding. Previous works have shown that mechanical instabilities play a crucial role in the formation of brain folds, and that the geometry of the fetal brain is one of the main factors in dictating the folding characteristics. However, modeling higher-order folding, one of the main characteristics of the highly gyrencephalic brain, has not been fully tackled. The effects of thickness inhomogeneity in the gyrogenesis of the mammalian brain are studied *in silico*. Finite-element simulations of rectangular slabs are performed. The slabs are divided into two distinct regions, where the outer layer mimics the gray matter, and the inner layer the underlying white matter. Differential growth is introduced by growing the top layer tangentially, while keeping the underlying layer untouched. The brain tissue is modeled as a neo-Hookean hyperelastic material. Simulations are performed with both, homogeneous and inhomogeneous cortical thickness. The homogeneous cortex is shown to fold into a single wavelength, as is common for bilayered materials, while the inhomogeneous cortex folds into more complex conformations. In the early stages of development of the inhomogeneous cortex, structures reminiscent of the deep sulci in the brain are obtained. As the cortex continues to develop, secondary undulations, which are shallower and more variable than the structures obtained in earlier gyrification stage emerge, reproducing well-known characteristics of higher-order folding in the mammalian, and particularly the human, brain.

## 1. Introduction

One of the most striking features of the human brain is its highly folded structure. Indeed, neuroscientists have for a long time pondered about its importance and origin (Bischoff, 1868; Cunningham, 1890). However, the process by which folds form is not yet fully understood, neither as a mechanical (Bayly et al., 2014) nor as a molecular process (Sun and Hevner, 2014). One of the main hypotheses to explain the convoluted nature of the cortex, commonly called the *differential tangential growth hypothesis* (Richman et al., 1975) posits that the buckling of the brain is created by a mismatch of growth rates in the cortical plate and the white matter substrate. The main contention with this hypothesis, however, is its requirement of a large difference between the stiffness of the two regions (Bayly et al., 2014). In order to obtain the wavelengths compatible with the gyral width of the human brain, the initial form of the differential growth hypotheses requires the stiffness ratio between the gray and white matter to be in the order of 10 (Richman et al., 1975). This is a major hurdle, as currently there is no consensus if the gray matter is indeed stiffer than the white matter, and if so, by how much. There is a solid body of evidence supporting the two possibilities, i.e., that the gray matter is indeed stiffer than the white matter (Budday et al., 2015; Green et al., 2008; Johnson et al., 2013), and vice-versa (Manduca et al., 2001; McCracken et al., 2005; van Dommelen et al., 2010). A second issue with the differential growth hypotheses is the shape of the sulci. This model results in smooth sinusoidal patterns, while the brain is characterized by smooth gyri and cusped sulcii (Tallinen et al., 2014).

The study of layered systems has been extensively conducted in the field of engineering, where it was used to model the buckling of sandwich-type panels (Hoff and Mautner, 1945), the Earth’s crust (Biot, 1957; Ramberg, 1970), etc. These works, however, deal mostly with stiff materials and large stiffness ratios. In recent years, there have been a surge in the number of works dealing with soft materials, with special focus in bio-compatible applications (Budday et al., 2017; Vandeparre et al., 2010) which provides an important tool in the understanding of the role of mechanics in the folding of the mammalian cortex.

Much work has been done to solve the issues with differential growth, specially on the sulci formation. For instance, Tallinen et al. (2014, 2016) performed large simulations to understand how the geometry and constraints affect the cortical folding, where they showed that the size and shape of the folds are dictated by the geometry of the early fetal brain. Other hypotheses have also been proposed to explain cortical gyrification. Perhaps best known in the medical community, van Essen (1997) conjectured that axonal traction is the driver of folding. Reaction-diffusion models, where the concentration and diffusion of growth-activator chemicals are explicitly modeled, have also been suggested as a way to explain both the gyrogenesis process, as well as the growth profile itself (Hinz et al., 2019; Verner and Garikipati, 2018).

Despite all these efforts, an aspect of brain folding that still remains elusive is the phenomenon of hierarchical folding, an important feature of the brain development, which must be included in order to understand the driving forces behind the complex folding patterns observed in the human brain.

Recent studies have analyzed the influence of growth and stiffness inhomogeneities along the cortex. A few years ago, Toro et al. performed studies on the effect of cortical inhomogeneity and curvature (Toro and Burnod, 2005). More recently, Budday et al. (Budday et al., 2015; Budday and Steinmann, 2018) performed similar inhomogeneity studies on rectangular geometries. In these works, structures with resemblance to the higher-order folding were obtained, but they lacked the complex spectrum of folding present in the human brain.

It has been shown that the thickness of cortex of the brain impacts the width and structure of brain folds (Armstrong et al., 1995; Budday et al., 2014; Heuer and Toro, 2019; Mota and Herculano-Houzel, 2015; Richman et al., 1975; Toro and Burnod, 2005). However, evidence on how competition between the different thicknesses in the cortex affect folding has been lacking. In this paper, the differential tangential growth hypothesis is augmented with an inhomogeneous cortical thickness field, yielding realistic folded structures, which could help explain formation of the deep sulci in the mammalian brain, hierarchical folding, as well as its consistent localization.

## 2. Material and Methods

We analyze two-dimension systems composed of two layers: a purely elastic, non-growing, softer substrate in the lower region, mimicking white matter, and a growing stiffer region on the top, emulating the cortical gray matter (see Fig. 1). The simulations are performed using a custom written finite element Method code to solve the continuum mechanics equations^1^.

**Figure 1:**
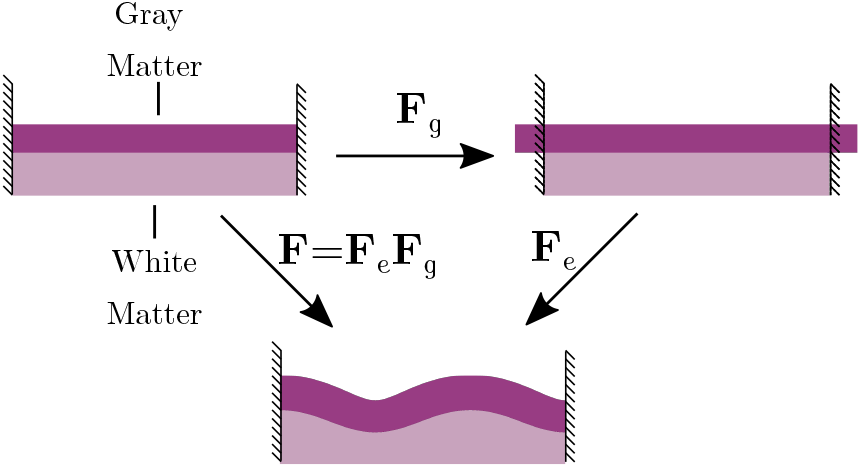
(Color online) Schematic representation of the model. The purple layer atop mimics the gray matter and is grown tangentially, while the pink substrate underneath mimics the white matter and does not grow. Growth is mathematically represented by the growth tensor **F_*g*_**, which can be discontinuous. In order to keep the compatibility with the attachment constraints between the gray and white matter, the system is subject to residual stress, described in this framework by the **F_*e*_** tensor.

### 2.1. Theoretical background

Due to the large-strain, nonlinear nature of the human brain, the framework of continuum mechanics (Bonet et al., 2016) is used. In order to distinguish between the original and deformed configurations, the following notation is introduced: The vector **X** denotes the coordinates of the original configuration, **x** the coordinates of the deformed configuration, and **u** = **x** − **X** denotes the displacement field. The deformation gradient tensor is written as

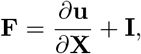

where **I** is the identity matrix. Growth is introduced using Rodriguez theoretical framework (Rodriguez et al., 1994), where the deformation gradient tensor **F** is decomposed into **F** = **F**_*e*_**F**_*g*_ (see Fig. 1), where, **F**_*e*_ describes the elastic part of the deformation and **F**_*g*_ the growth contribution. The energy and stress are calculated from the elastic part of the deformation gradient tensor alone. Thus, in this framework, the energy-density (and all quantities derived) is defined in terms of **F**_*e*_.

The soft tissue of the brain is modeled by the compressible Neo-Hookean energy-density function (Bonet et al., 2016)

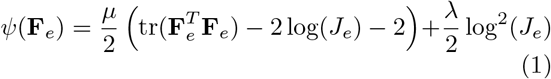

where *J_e_* = det(**F**_*e*_), and *μ* and λ are the Lamé parameters. This energy-density family has been shown to appropriately model the brain tissue (Budday et al., 2015, 2017).

Due to the relatively long time scale of cortical development when compared to the elastic response of brain tissue, the quasi-static approximation is used. At every value of **F**_*g*_ the displacement field **u** is calculated, obeying the equilibrium equation

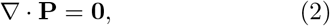

where **P** is the first Piola-Kichhoff stress tensor, related to the energy-density *ψ* by

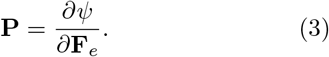

Both, Eq. 2 and Eq. 3 retrieve the functional form of their classical continuum mechanics counterparts in the limits of no growth, i.e., **F**_*g*_ = **I**. The values of λ and *μ* are chosen such that in the linear (i.e., small strain) regime, the Young moduli ratio between the gray matter (GM) and the white matter (WM) is *E_GM_/E_WM_* = 3, consistent with previous models (Budday et al., 2015) and the Poisson ratio *ν* = 0.35 on both layers. Due to the nonlinear nature of Eq. 1, the value of the Poisson ratio *ν* and Young modulus are dependent on the current displacement in the system. Specifically, they depend on the determinant *J_e_* (Bonet et al., 2016) as

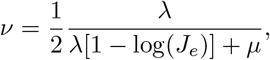

and

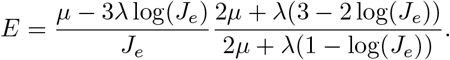

Notably, the energy-density in Eq. 1 has no inherent length scale. Thus, only the ratios between the elastic moduli are important for the phenomenology presented in this paper.

### 2.2. Simulation details

The cortical layer is grown linearly, i.e., the growth tensor is described by

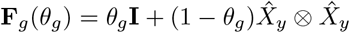

where the growth parameter *θ_g_* measures the degree of elongation in the cortex. For instance, at *θ_g_* = 2 the gray matter would have expanded to twice its lateral size if it were not constrained. To mimic differential growth, *θ_g_* is varied in the interval [1.0, 2.5] in the gray matter region, while it is kept at unity in the white matter region. The growth parameter *θ_g_* is increased in small steps of 0.01.

At every growth step, Eq. 2 is solved using the finite element method, with boundary conditions of zero displacement on the bottom surface *X_y_* = 0 and zero stress on the top surface *X_y_ = L_b_*. In order to minimize boundary effects, periodic boundary conditions are imposed on the sides of the surface, *X_x_* = 0 and *X_x_ = L_b_*. The box lengths *L_b_* will be specified in each section. The system being two dimensional, corresponds to an infinite system in the *z*-axis, with the constraint of no displacements in the *z*-axis.

Due to the nonlinear nature of the energy described in Eq. 1, the divergence of the first Piola-Kichhoff stress tensor will also be nonlinear. To find the roots of this function, the Newton method augmented by a backtracking algorithm (Bonet et al., 2016; Nocedal and Wright, 2006) is used. In order to avoid overlaps, collisions are detected and resolved using the approach introduced by McAdams et al. (2011). As any collisions will be initiated in the cortical region, calculations are optimized by only detecting collisions in the gray area elements. This generates no artifacts, as due to the structure of the mesh, collisions are initiated in the gray area, and are resolved before any white matter elements are involved.

In order to allow the system to overcome metastable states, a small force field pointing in the *X_y_* direction is introduced. The forces are drawn from a uniform random distribution between [−7 × 10^−2^, 7 × 10^−2^)×*E_GM_*. Each simulation has been repeated three times with different seeds. No significant difference is observed between runs with different seeds, or when the forces are doubled or halved.

## 3. Results

### 3.1. Homogeneous Thickness of Cortical Ribbon

We analyze the folding of a slab with constant cortical thickness *T* throughout, in the range [0.1, 0.5]cm. In this case, it is expected that the system will fold into well defined wavelengths, with no localization (Groenewold, 2001). In order to avoid finite-size effects, the simulation box is set to *L_b_* = 100cm ≫ λ_*F*_, where λ_*F*_ is the folding wavelength.

Initially (i.e., for little growth) even though the growth happens tangentially, the cortex does not increase in length, but rather in thickness due to the confinement and resulting stress (see Fig. 2). Eventually, at a critical growth 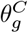, the stress exceeds the critical buckling threshold, and the system buckles in a well defined, almost sinusoidal wave pattern (see Fig. 3). As growth continues, the wavelength of the pattern and the average thickness remain almost unchanged, while the amplitude increases. Interestingly, the thickness of the cortex is no longer homogeneous. Sulci (regions of positive curvature) are significantly thinner than gyri (regions of negative curvature). As growth continues post buckling, the difference in thickness continues to increase.

**Figure 2:**
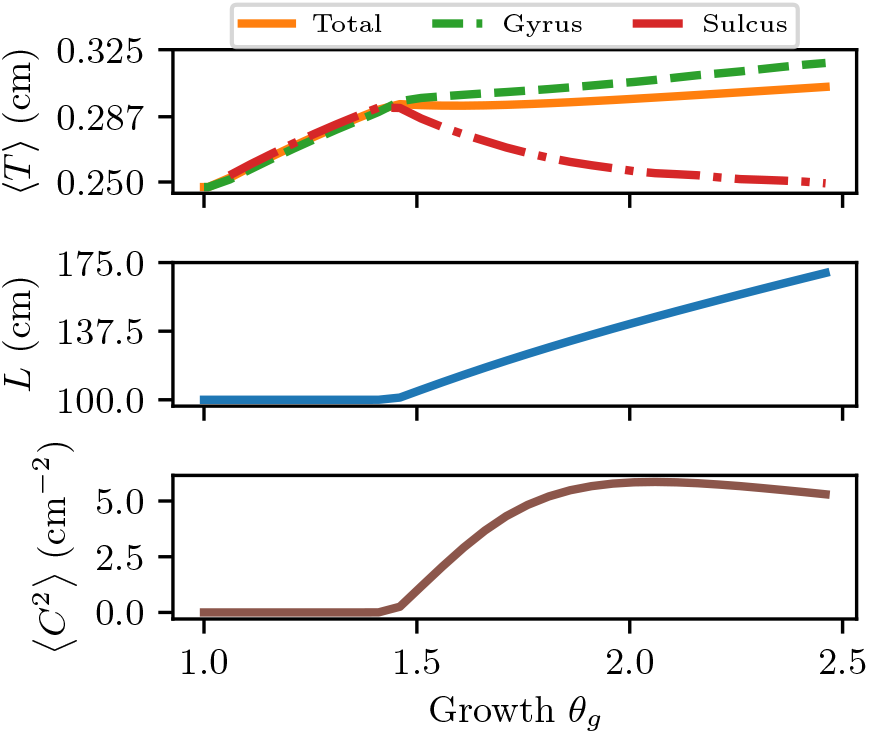
(Color online) Spatial averages of several observables in the system with initial thickness *T* = 0.250 cm, as a function of the growth parameter *θ_g_*. In (a), the dependence of the average thickness with growth are shown, as well as the average cortical thickness of the sulci and the gyri, as defined in the text. In (b) and (c) the contour length of the top layer of the system and squared curvature of the system are shown, respectively. This last quantity is specially useful to characterize the onset of folding, as will become clear in Sec. 3.2.

**Figure 3:**
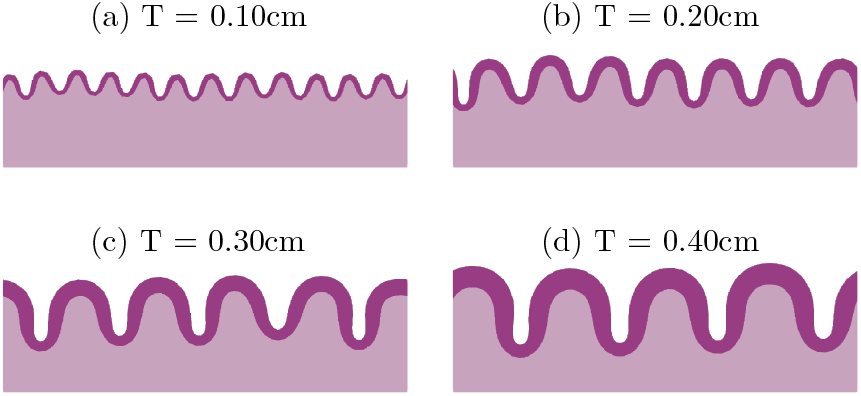
(Color online) Folded system for the thickness indicated within the figure. In order to improve visualization, only the region *y* > 95cm is shown here.

In each simulation the Fourier transform of the function *u_y_*(*X_x_*) (i.e., the displacement in the *y* direction as a function of the material coordinates) is calculated along the top layer of the system. The weighted average wavenumber and wavelength are then obtained as

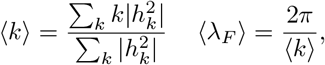

where *h_k_* is the coefficient of Fourier expansion for the mode with wavenumber *k*. In order to obtain the weighted average wavelength for each cortical thickness *T*, the simulations are repeated with inceasing number of mesh cells, ranging from 2^12^ to 2^20^ cells. The value of the weighted average wavelength is then obtained by wavelength via a linear extrapolation to the infinitely refined mesh.

The weighted average wavelength increases linearly with initial thickness *T* (see Fig. 4). The linear dependency can be obtained from simple dimensional analysis: As the elastic equations have no inherent length scale, the cortical thickness is the only relevant length scale, as long as system size is not limiting. Thus, changing the homogeneous cortical thickness can be seen as a change of measurement units, or progressive zooming in on the same base system (see Fig. 5). It is possible to estimate the slope for a linear elastic substrate through analytical calculations as (Groenewold, 2001)

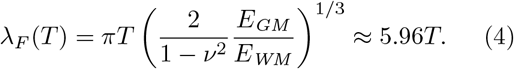

**Figure 4:**
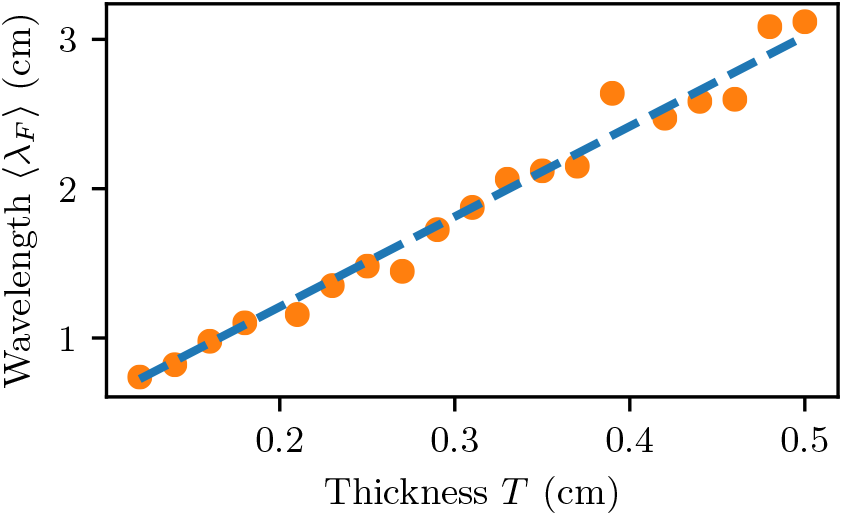
(Colors Online) Weighted wavelength of the system as function of the cortical thickness *T*. The orange circles indicate extrapolation results, while the broken blue lines represent the linear fit 〈λ_*F*_〉 = 6.04*T* cm with *R*^2^ = 0.98.

**Figure 5:**
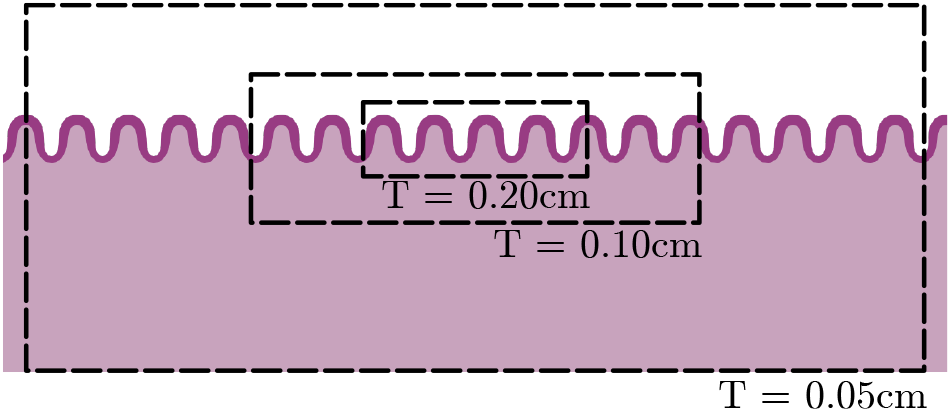
(Color online) Schematic representation of scaling of the system. Due to the lack of inherent length scale in the elastic equations, systems with thicker cortices can be seen as subregions of systems with thinner cortices. Here, each broken rectangle highlights a region which is equivalent to a system with the cortical thickness *T* indicated within the figure.

Such slope presents a weak dependence on the stiffness ratios between the gray and white matter, obeying a weak power law. Thus, the specific value of the ratio *E_GM_/E_WM_* plays only a minor role in the determination of the folding wavelength.

### 3.2. Inhomogeneous Thickness of Cortical Ribbon

The cortical thickness of the brain is spatially inhomogeneous. In order to emulate this inhomogeneity, a variable cortical thickness *T*(*X_x_*) is introduced. Specifically, as a generic form of thickness variation, a sinusoidal thickness variation of the form

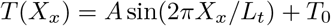

is chosen. This inhomogeneity introduces two new length scales beyond the base thickness *T*_0_: the inhomogeneity amplitude *A* and the period of the inhomogeneity *L_t_*. Thus, in contrast to the previous results, it is possible to choose any one of the three as the fixed length scale, and vary the remaining two independently. In this study, the thickness period *L_t_* = 10 cm is chosen as the fixed scale. The folding pattern for a different periodicity 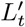 can be obtaining by rescaling the spatial quantities by 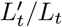, or a suitable power thereof.

Note that any form of thickness variation can be written as a sum of sinusoidal variations. When deformations are small, even the resulting folding patterns can be obtained by simple superposition. In the brain, however, deformations are large and nonlinear, and each thickness field must be studied independently.

Simulations are performed for base thickness in the same range as before, [0.1, 0.5] cm, and for each *T*_0_, the amplitude *A* was varied in the range [0, 0.9] × *T*_0_. The inhomogeneity creates a much more localized deformation, thereby reducing possible artifacts created by finite-size effects. Thus, in order to maximize computational efficiency, the simulation box is chosen as *L_b_* = *L_t_*.

If the natural folding wavelength λ_*F*_ of the local thickness is much smaller than the periodicity length, the system behaves essentially like the homogeneous systems studied in Section 3.1, i.e., the folding wavelength obeys Eq. 4, with *T* = *T*(*X_x_*). Here, the system folds into well defined waves, but with the spatially dependent wavelength commensurate with the cortical thickness of the underlying region (see Fig. 6 (a), (c)). Accordingly, these systems also present the constant length - constant thickness stress-relief mechanism.

**Figure 6:**
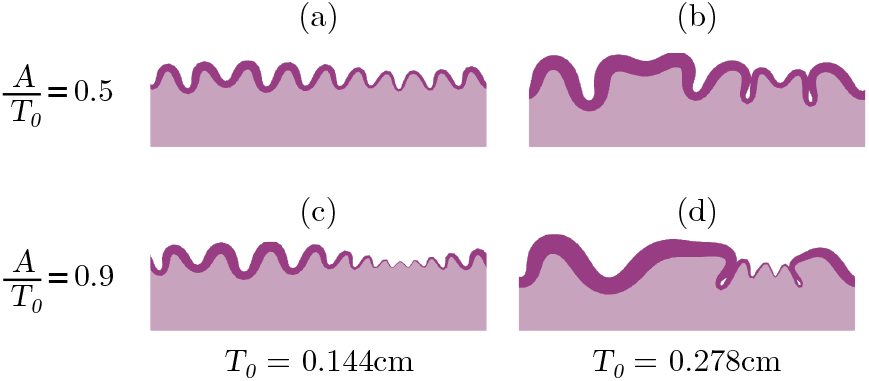
(Color online) Simulations with varying inhomogeneities amplitudes *A* at growth parameter *θ_g_* = 2.5. The simulations have a base thickness *T*_0_ as indicated within the figure.

However, when the folding wavelength becomes comparable with the periodicity length, a second form of folding arises, characterized by complex folding patterns. In these conformations, several wavemodes are simultaneously obtained (see Fig. 6 (b), (d)), presenting similarities with the further regions of the gyrencephalic brain.

This new shape has distinct developmental steps, which differ from those described in Sec. 3.1. For small growth, a initially flat system (see Fig. 7 (a)) forms a single, deep, sulcus in an otherwise planar cortex, in the region surrounding the thickness minimum (see Fig. 7 (b)). The depth of this sulcus soon saturates, and due to the underlying white matter, it is energetically favorable to form additional sulci rather than to increase the depth of the exist sulcus as growth continues (see Fig. 7 (c)). The maturation of the new sulci occur concurrently with the formation of shallow sulci in the regions of highest thickness.(see Fig. 7 (d))

**Figure 7:**
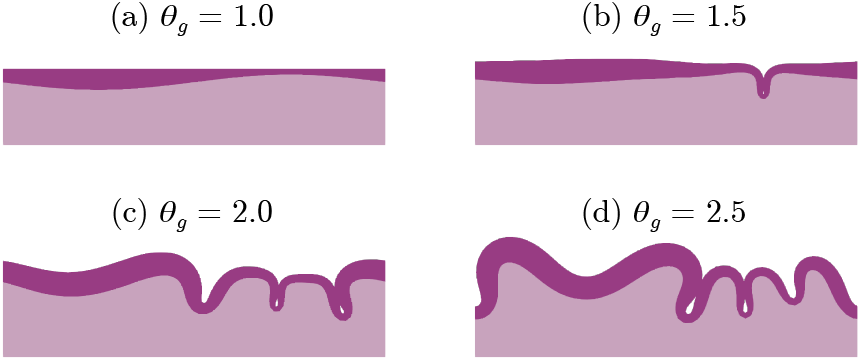
(Color online) Growth evolution of system with *T*_0_ = 0.45cm and *A* = 0.315 cm. The growth parameters are indicated within the figure.

It is expected that the folding starts on the region of thinnest cortex. In the limit of small deformations, the system can be analysed by the theory of thin plates. In this domain, the bending rigidity depends on the cube power of the thickness of the plate (Ventsel and Krauthammer, 2001). Thus, the large differences in thickness create a stress imbalance in the region, leading to the buckling of the region with small thickness. This is specially noticeable in the formation of the deep sulci observed in Fig. 7. Here, the thick parts of the cortex compress laterally, which leads to stress condensation in the thin parts of the cortex. The thin region has then to absorb the compression of the whole system.

The simulations are in qualitative agreement with the results from linear stability analysis. The details how this theory is applied to our model are outlined in Appendix A. In short, the lowest-energy mode of a bilayer system is calculated, where we introduce a thin plate as the top layer, and an elastic substrate underneath. The upper plate has a sinusoidally varying bending rigidity

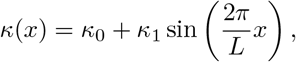

where *L* is the periodicity length. The system is then subject to a spatially constant compressive surface tension *γ*. The bending rigidity plays a similar role in this model as the thickness plays in the simulations, and the surface tension plays a similar role to growth.

According to the analytical model, homogeneous systems (i.e., *κ*_1_ = 0) will fold into a single, well-defined wavenumber, as expected (see Fig. 8 (a)) (Hornung, 2019). However, as the bending rigidity ratio *κ*_1_/*κ*_0_ is increased, the folding gets more localized around the bending rigidity minimum. This phenomenon is qualitatively consistent with what was observed in the simulations. For the case with *κ*_1_/*κ*_0_ = 0.9, for instance, the linear stability analysis predicts the formation of a deep sulcus surrounding the bending rigidity minimum, similarly to the finite-element simulations (see Fig. 7 (b)). The analytical model predicts that as the cortical plate gets more compressed, the system develops secondary sulci, as can be noticed in region *x/L* ≈ 0.25 in the system with *γ* = −1000*κ*_0_/*L* (see Fig. 8 (b))

**Figure 8:**
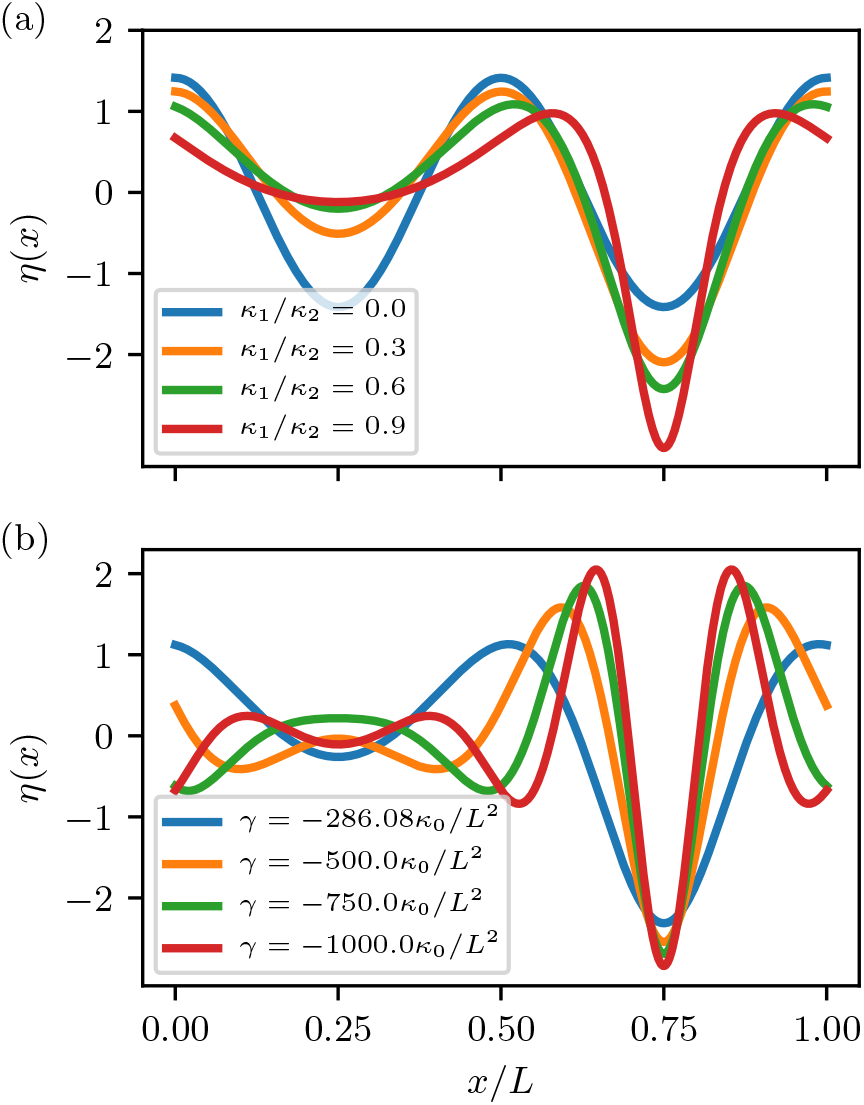
(Color online) Analytical prediction of vertical dislocations *η*. (a) Varying bending rigidity ratios at buckling point. (b) Varying values of surface tension *γ*, while the bending rigidity ratio is kept constant as *κ*_1_/*κ*_0_ = 0.5. For *γ* > − 286.08*κ*_0_/*L*, no buckling is predicted. Note the appearance of higher order folding for *γ* < − 1000*κ/L*. In both cases, the effective Young Modulus used was 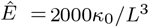. As the curves are normalized, the predicted dislocations can only be compared within the same curve, but not between curves calculated with distinct parameters.

It is possible to compare the structures obtained to histological sections of the human brain, as shown in Fig. 9. The results from our simulations present a striking similarity to some regions of the human cortex, showing a wide range of sulcal depths and widths, in qualitative agreement with the ones observed in regions with higher-order folding. For instance, the superior parietal lobe presents a plethora of small, shallow folds, similar to those observed in the simulations with thin cortices. Regions presenting a more complex folding pattern, such as the postcentral gyrus, or the posterior middle temporal gyrus are reproduced by simulations with thicker cortices. Furthermore, the gyral height-to-width ratio resulting from the simulations are similar to those observed in the histological sections.

**Figure 9:**
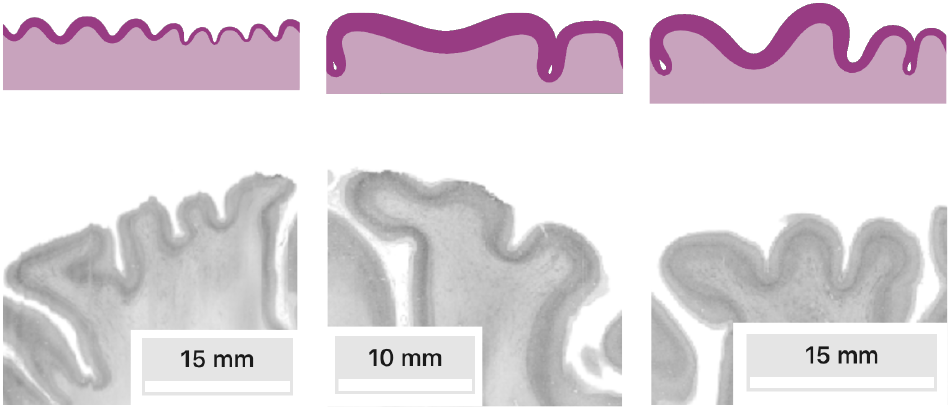
Illustrative comparison between simulation results (top) and sections of the cortex (bottom; adapted from HBP BigBrain (Amunts et al., 2013)). From left to right, the simulations are performed with *T*_0_ = 0.189 cm, *A* = 0.102 cm; *T*_0_ = 0.500 cm, *A* = 0.180 cm; *T*_0_ = 0.367 cm, *A* = 0.198 cm. In the same order, extracts of the left superior parietal lobule (sagittal plane), the right postcentral gyrus (coronal), and the right posterior middle temporal gyrus (coronal) are shown. Due to the arbitrary choice in the value of *L_t_*, the thicknesses between the simulation and the histological section are not quantitatively comparable.

Next, we turn our attention to the onset of buckling, i.e. the critical amount of growth 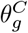 above which the system starts to fold, broadly indicating when the constant-length regime ceases, and the constant-thickness starts. The critical growth 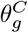 is defined somewhat arbitrarily as the growth where the averaged squared curvature of the system reaches 2cm^−2^. The results do not change considerably for choices of critical curvature square 〈*C*_2_〉 in the range [1, 3]cm^−2^. The transition points are shown in Fig. 10, where it is possible to observe that the critical growth 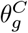 is strongly affected by interplay between the inhomogeneity amplitude and the base thickness of the cortical plate, with a noted decrease in the value of 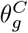 as the inhomogeneity amplitude *A* increases. In light of these results and of the differences between the folding of thin and thick systems (see Fig. 6), we conjecture that the relevant parameter to cortical convolution is not solely the ratio between the maximal and minimal thickness, but that the local gradient of the cortical thickness also plays a fundamental role.

**Figure 10:**
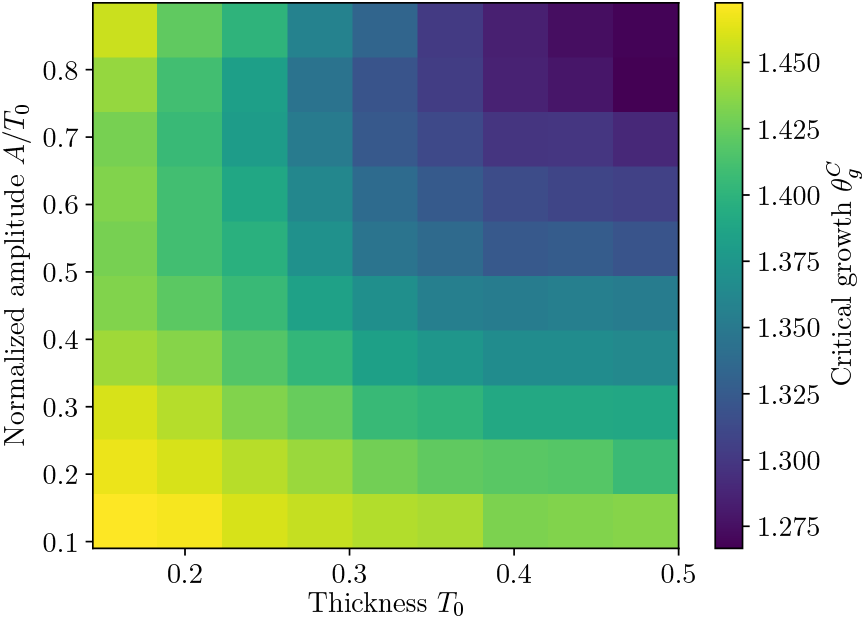
(Color online) Critical growth 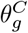 for the emergence of folding for various combinations of cortical thickness and inhomogeneity amplitudes.

## 4. Conclusion

We have analyzed the effects of the cortical inhomogeneity in the formation of brain folds. To this end, two closely related systems were studied. First, analyses was carried out by simulating a rectangular bilayer slab where the top layer grows tangentially. It was observed that the folding pattern follows well-defined wavelengths, which depended on the thickness of the top layer, consistent with previous work (Biot, 1937; Budday et al., 2014; Groenewold, 2001). According to Bok’s principle, the thickness differences between the sulci and gyri are created as a consequence of the curvature of the brain. During gyrogenesis, the high curvature in the sulci spreads the cortical mantle, decreasing its thickness. The gyral crowns are not affected as strongly due to their relatively small curvature (Bok, 1929). Our simulations are consistent with this principle, showing that an initially homogeneous cortex can develop cortical inhomogeneities through buckling, coherent with prior observations on homogeneous systems (Holland et al., 2018; Riccobelli and Bevilacqua, 2020; Welker, 1990).

Second, the effects of wavelength competition were studied through the introduction of inhomogeneities in the cortical thickness. In these systems, phenomena closely related to the mammalian gyrogenesis, such as the emergence of hierarchical folding in systems with thick cortices, were observed. Indeed, the results shown here indicate that inhomogeneities in the cortical thickness might play an important role in the localization and formation of hierarchical folding patterns of the brain. Specifically, it was shown that these inhomogeneities are sufficient to break the simple wave-like patterns observed in the homogeneous system. Further, our observations indicate that thickness inhomogeneity leads to earlier folding compared to systems with homogeneous cortical thickness. Lastly, the results obtained *in silico* were proved to be consistent with those obtained from analytical models derived from thin plate theory. Similar approaches have been taken in other studies, where other cortical inhomogeneities were studied both in circular (Toro and Burnod, 2005) and rectangular (Budday and Steinmann, 2018) geometries. Our results are in line with those findings, but exhibiting a more complex folding pattern, as well as the emergence of multiple sulci within each inhomogeneity period.

The computational model used to explain the hierarchical folding patterns of the mammalian brain is very general, utilizing only the fundamental elastic nature of brain tissue. The absolute values of the elastic moduli play no role, with only their ratios being important. This model is agnostic to all sorts of structural properties of the brain, such as its volume, eccentricity, functional connectivity, etc. The results derived here are thus applicable to the brains of other mammalians, after a suitable scaling of the thickness, growth, and periodicity lengths.

While the comparisons presented here focus mostly on smaller regions of the human brain, we conjecture that more extreme forms of inhomogeneity can lead to the formation of the deep sulci obtained in the mature human brain. Larger simulations are required to gauge the influence of thickness inhomogeneity on the full brain. Based on our current results, the mutual influence of the different kinds of inhomogeneity (e.g., thickness and growth) can also be studied.

We have shown the consequences of cortical thickness inhomogeneities to brain folding. What drives the development of these inhomogeneities is still a matter of ongoing research. Features of a given brain area, such as its cortical thickness are related to its function (Geyer and Turner, 2013). Biomolecular and mechanical factors contribute to that (Kriegstein et al., 2006; Sun and Hevner, 2014), and, based on our current approach, could additionally be included in future models of brain folding.

It can also be shown that viscoelastic properties can affect the buckling wavelength of bilayered systems (Biot, 1957). Thus, it would be interesting to further refine the current model with the viscoelastic characteristics of growing tissues in general (Ranft et al., 2010), and of the brain in particular (Budday et al., 2017). Finally, structural connectivity, i.e., axons within the white-matter fiber tracts, have been conjectured to be one of the drivers of folding (van Essen, 1997). How they would influence the conformations found in this work could be investigated further to understand the mutual influence of differential growth and axonal tension in gyrogenesis.

This paper’s focus was the developing brain, but in virtue of generality of the model used, its results are extensible to other fields. For instance, it has been shown that microscopic corrugated surfaces give rise hyperhydrophobic surfaces, in the so called Lotus effect (Gao and McCarthy, 2006; Marmur, 2004). Our results can provide further insight into the self assembly of these corrugations, and in the control of their properties.

Soft layered systems can be realized experimentally by gel slabs coated with gels with different properties (Auguste et al., 2014; Budday et al., 2017). These systems have been used as simulacra for brain folding (Holland et al., 2018; Tallinen et al., 2016), where they were able to mimic the folding a 3D-printed human brain. Thus, the production of samples with sinusoidal variation of the top-layer thickness would allow for the experimental testing of our predictions.

## 5. Acknowledgments

The authors gratefully acknowledge the computing time granted by the JARA-HPC Vergabegremium on the supercomputer JU-RECA (Jülich Supercomputing Centre, 2018) at Forschungszentrum Jülich. Simulations were additionally performed with computing resources granted by RWTH Aachen University under project rwth0399. We would like to thank Dr. Claude J. Bajada for helpful discussions and feedback. This work was further supported by a grant from the Initiative and Networking Fund of the Helmholtz Association (SC) as well as the European Unions’s Horizon 2020 Research and Innovation Program under Grant Agreement 785907 (Human Brain Project SGA2; SC). LCC also acknowledges the support by the International Helmholtz Research School of Biophysics and Soft Matter (IHRS BioSoft).

## Appendix A. Simplified Analytical Model

In order to understand the buckling of the system, we use a simplified model which can be solved analytically. Here, a thin plate with a spatially varying bending rigidity is studied. This plate is attached to a linear elastic substrate, filling the whole of the half-space *y* < 0. In the limit of small deflections and disregarding shearing, it is possible to write the displacement of the system in the Monge representation as

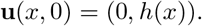

Here, *h*(*x*) indicates the local height of the plate along the *x* axis. The free energy of this system is composed of three terms:

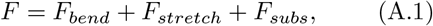

where *F_bend_* is the free energy of bending the thin plate, *F_stretch_* is the energy required in order to stretch the plate and *F_subs_* is the energy of the deformed underlying substrate. Explicitly,

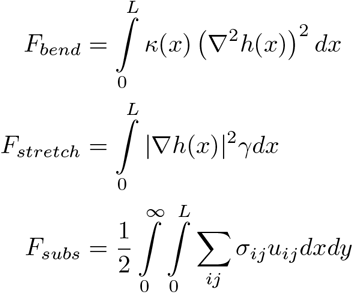

where *κ*(*x*) describes the space-dependent bending rigidity, *γ* is the surface tension on the superficial plate, *σ_ij_* are the components of the Cauchy stress tensor, and *u_ij_* are the components of the strain tensor 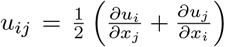 In order to keep the simplicity of the model the bending rigidity of the plate, rather than its thickness, varies sinusoidally. That is,

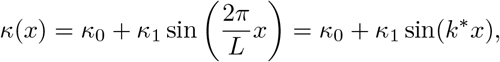

where *k*\* is the characterstic wavenumber of the inhomogeneity. Due to the periodic nature of our system, the stability analysis is easier to carry out in Fourier space. Thus, the height function *h*(*x*) is expanded into

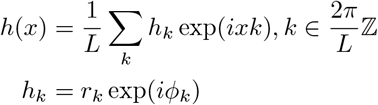

with *ϕ_k_* = *ϕ_−k_*. In this decomposition, whole-plate dislocation(i.e., *h*_0_ = 0) were disregarded.

To obtain the energy of the elastic substrate in the Fourier space, one has to solve the problem of a linear elastic substrate with given surface deformation in Fourier space, as derived in Ref. (Groenewold, 2001). With this solution, the free energy described in Eq. A.1 is written as

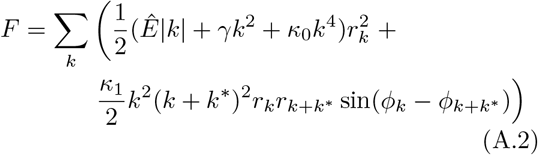

with

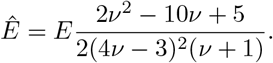

We search for the buckling modes that are most unstable. That is, those with wavenumber *k* which minimize the energy in Eq. A.2. Due to the bound properties of the sinus, it is clear that the condition *ϕ_k_ − ϕ_k+k*_* = 3/2*π* + 2*πn* is necessary to obtain this minimum. Thus, the energy is written as

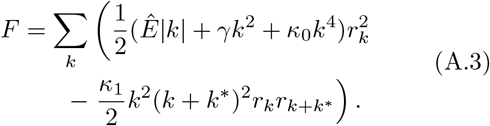

Unstable modes can then be obtained by standard stability analyses. Eq. A.3 is recast into its matricial form

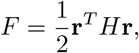

where **r** is a vector with components **r** = *r_k_*, and *H* is the Hessian matrix. Explicitly,

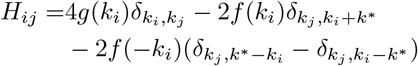

with

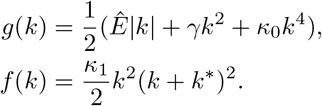

In this form, the unstable modes are obtained as those states with negative eigenvalues for the Hessian matrix, corresponding to modes with negative energy. For homogeneous systems (i.e., *κ*_1_ = 0), the energy contribution of each mode is independent (i.e., the Hessian matrix is diagonal), and modes that minimize the energy can be obtained analytically (Hannezo et al., 2011; Hornung, 2019). In the inhomogeneous case, the various wavemodes are coupled, and it is necessary to solve the eigenproblem numerically. Fig. 8 shows the results of these calculations. Each curve corresponds to the eigenfunctions with lowest eigenvalues for different elastic parameters, as indicated therein.

This analytical theory gives results which are qualitatively similar to those obtained in our simulations. In order to obtain quantitative comparisons, a more complex theory is necessary, which takes into account the lateral displacements during gyrification.

## Footnotes

1 The simulations were written using the deal.II library (Alzetta et al., 2018; Bangerth et al., 2007), and parallelized using MPI via PETSc (Balay et al., 2019a,b).

